# Particle size distribution shift and diurnal concentration changes of environmental DNA caused by fish spawning behaviour

**DOI:** 10.1101/2023.06.19.545639

**Authors:** Satsuki Tsuji, Naoki Shibata

## Abstract

Spawning is one of the most significant aspects of fish life history, so understanding spawning is critical for the conservation and management of species. Recently, an efficient and non-invasive spawning detection method for spawning behaviour has been proposed, utilising the increased environmental DNA (eDNA) concentration and nuclear/mitochondrial eDNA ratio associated with fish spawning. However, little is known about the characteristics and dynamics of sperm-derived eDNA, which is key to detection. This study focused on changes in eDNA particle size distribution (PSD) and concentrations pre-, post-, and during spawning of Ayu, *Plecoglossus altivelis*. Firstly, PSD changes between pre- and post-spawning were investigated by comparing concentrations and proportions of eDNA obtained from filters with different pore sizes. Secondly, the diurnal changes in eDNA concentration were monitored at the peak of the spawning season by collecting river water every one or three hours. Results showed that eDNA related to sperm-head size increased at post-spawning, and eDNA concentrations had significant diurnal changes with a peak during the spawning time window. These findings suggest that semi-selective recovery of sperm-derived eDNA based on particle size and/or sampling during the spawning time window with increased concentrations can improve the detection sensitivity of eDNA-based spawning surveys. This study provides essential basic information for advancing eDNA-based spawning surveys and contributes to their further development.

## Introduction

The utilization of environmental DNA (eDNA) analysis for the evaluation and administration of aquatic ecosystems is rapidly expanding and now becoming commonplace (Bohmann et al., 2014; Lodge et al., 2012; Rees et al., 2014; Taberlet et al., 2012; Thomsen & Willerslev, 2015; Tsuji et al., 2019). Environmental DNA analysis enables us to indirectly reveal the presence and/or biomass of macroorganisms based on the detection of DNA materials derived from their metabolic waste, sloughed skin cells and gametes released into the environments including soil, water, and air (Bylemans et al., 2017; Ficetola et al., 2008; Kelly et al., 2014; Kuwae et al., 2020; Lynggaard et al., 2022). Moreover, recent studies have shown that the eDNA concentration and the nuclear DNA (nuDNA)/mitochondrial DNA (mtDNA) ratio may increase during the spawning season compared to the non-spawning season (Bayer et al., 2019; Ip et al., 2023; Spear et al., 2015; Takahashi et al., 2018; Thalinger et al., 2019; Tillotson et al., 2018). This phenomenon is attributable to the fact that gametes, primarily sperm, discharged during spawning behaviour, are detected as eDNA in species that perform external fertilisation (Bylemans et al., 2017; Tsuji & Shibata, 2021).

By focusing on the eDNA concentration and/or ratio alterations that are distinct to the non-spawning season, eDNA analysis has the potential to enable us to estimate species spawning behaviour and its magnitude and to detect species with small biomass more sensitively. An understanding and knowledge of spawning are crucial for the conservation and management of species because it constitutes one of the most significant aspects of their life history (Danylchuk et al., 2011; Spear et al., 2015). Spawning surveys using eDNA analysis are potentially useful because they provide an opportunity to monitor and understand spawning with less effort, time, and invasiveness than conventional observation-based methods (Inui et al., 2021; Wu et al., 2023). Additionally, it is probable that appropriate sampling during the spawning season, when eDNA concentrations are temporarily heightened, will enhance the probability of detecting species with small biomass (Bracken et al., 2019; Crane et al., 2021). The low eDNA concentrations are one of the primary reasons for false-negative results in surveys focused on detecting endangered species and recently introduced non-native species (Carim et al., 2019; Jerde et al., 2013). A sampling strategy targeted at the spawning season could reduce the risk of false negatives.

Despite the key role of gamete-derived eDNA in eDNA studies during the spawning season, its characteristics and dynamics remain largely unexplored (Tsuji et al., 2022). As eDNA detectability and persistence are significantly influenced by its characteristics and dynamics, understanding these aspects is critical for designing effective sampling strategies and interpreting results accurately (Harrison et al., 2019). While substantial knowledge has been accrued on somatic-derived eDNA (such as from skin, feces, and mucus), it is pointed out that previous knowledge may not be applicable because the morphological features of gametes-derived eDNA differ significantly from them (Barnes & Turner, 2016; Tsuji et al., 2022; Ulloa-Rodriguez et al., 2017). Teleost spermatozoa, for example, the range is between about 25 and 100 μm in length and mainly consists of a head tightly overlayed with the plasma membrane, a middle section containing a mitochondrial capsule with multiple disulfide bridges, and a long tail (Bobe & Labbé, 2010; Ulloa-Rodriguez et al., 2017). These sperm are released into the water in an almost intact state, and the size of each part is almost uniform within each species.

This study focused on examining the changes in particle size distribution (PSD) of eDNA pre- and post-spawning, as well as the diurnal changes in eDNA concentration during the spawning season. Firstly, the understanding of PSD is crucial for estimating the dispersion and settling rate of eDNA, and it may allow for selective recovery of gamete-derived eDNA based on particle size. Selective recovery of sperm-derived eDNA could enhance the sensitivity of spawning activity detection, as well as aid in assessing genetic diversity within spawning groups. Secondly, since the increase in eDNA concentrations during the spawning season is mainly attributed to sperm release (Tsuji & Shibata, 2021), eDNA concentrations are likely to show diurnal changes with peaks at the spawning time window. In the only previous study, spatiotemporal changes in eDNA concentrations post-spawning were investigated using a single carp (*Cyprinus carpio*) pair in experimental ponds (Wu et al., 2022). Their data simulations found that concentrations peaked by spawning, distributed uniformly in the experimental pond, and returned to baseline within approximately 24 hours. Nevertheless, in natural lotic habitats, eDNA concentrations may change more dynamically and over a shorter period due to multiple spawning individuals and continuous diffusion. Therefore, if this assumption is correct, it will be essential to pay attention to the timing of water sampling for conducting an accurate and quantitative survey based on eDNA analysis to monitor spawning.

This study aimed to explore pre- and post-spawning changes in eDNA particle size distribution (PSD) and diurnal eDNA concentration shifts during the fish spawning season in a river. Two experiments were conducted in a river using Ayu (*Plecoglossus altivelis*), which is known to spawn daily in the hours post-sunset during their spawning season, as a model species. In experiment 1 (Exp. 1), we collected water samples pre- and post-spawning at the initial and peak phases of the spawning season, examining eDNA PSD for two genetic regions. In experiment 2 (Exp. 2), diurnal changes in eDNA concentration were monitored during peak phases of the spawning season by time-series sampling over a 27-hour period with emphasis on the spawning window. Based on the results, we discussed the potential of selective recovery of sperm-derived eDNA, its advantages and the need for careful consideration of sampling timing in view of spawning ecology.

## Materials and Methods

Overview of the experimental designs are presented in Fig.1. Both experiments were conducted in the lower reaches of Shiotu-o River, Japan (35°30’58.7”N 136°09’48.8”E, Fig. S1). The Shiotsu-o River flows into Lake Biwa.

**Figure 1.**
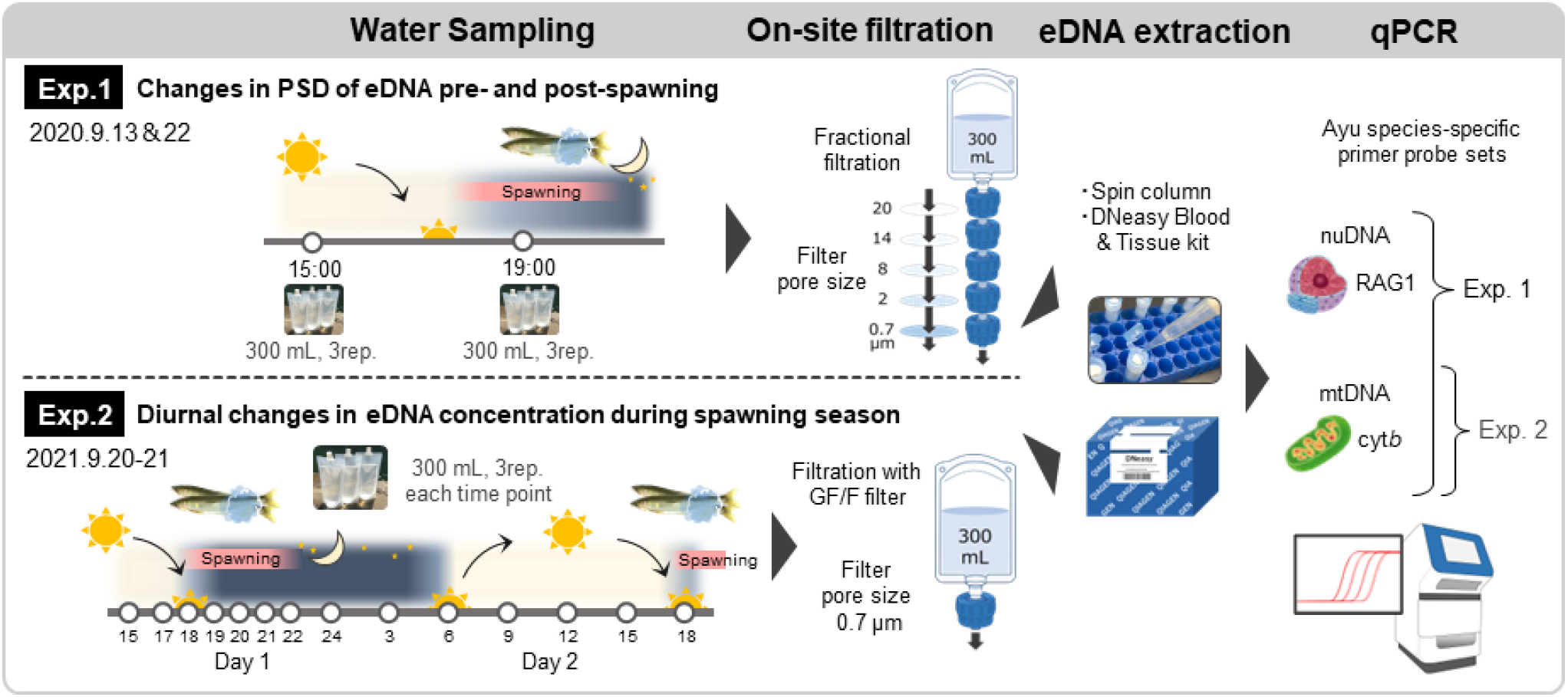
Overview of the two experimental designs

### Experimental species

Ayu (*Plecoglossus altivelis*) is an important species for fisheries and recreational fishing in Japan and a key member of Japanese riverine ecosystems because of its unique ecological position as an algivorous specialist (Iguchi et al., 2002; Kawanabe, 1996; Miyadi, 1960). The life cycle of Ayu is one year, with an amphidromous life history in which the larvae migrate from coastal areas to rivers in spring for further growth, maturation, reproduction and death (Azuma, 1973; Nishida, 1974). In Lake Biwa, the landlocked Ayu population inhabits and can be divided into two main groups with different life histories: the amphidromous type mentioned above and the anadromous-like type, which inhabits the lake until maturity and migrates to the river just before spawning (Azuma, 1973; Iguchi et al., 2002; Iguchi & Nishida, 2000). However, regardless of life history type, spawning occurs in the autumn in the lower reaches of rivers. During the peak of the spawning season of landlocked Ayu in Lake Biwa (roughly late September to middle October), they form dense colonies and spawn daily around sunset to several hours on gravel beds in riffles (Miyadi, 1960; Takahashi & Azuma, 2016). It means that eDNA derived from spawning, primarily sperm-derived eDNA, is only released within a limited time window. The sperm of Ayu is primarily composed of an oval capsule-shaped head (measuring 2.1 μm in length and 1.2 μm in width) and a lengthy flagellum (21.2 μm) (Fig. S2) (Hara & Okiyama, 1998; Ulloa-Rodriguez et al., 2017). The sperm head contains a nucleus and some mitochondria.

### Experiment 1: Changes in concentration and PSD of eDNA pre- and post-spawning

Water sampling was carried out on September 13th (the earliest phase of the spawning season) and 22nd (the peak phase) in 2020. The spawning status around each survey date was confirmed by direct egg observation by the Shiga Prefectural Fisheries Experiment Station (https://www.pref.shiga.lg.jp/file/attachment/5310978.pdf; Table S1). 300 mL of surface water was collected in triplicate at 15:00 (daytime, pre-spawning) and 19:00 (one hour after sunset, during spawning) using disposable bags (DP16-TN1000, Yanagi, Aichi, Japan). The collected water was fractionally filtered on-site via a series of filters with different pore sizes, 20, 14, 8, 2 μm (Track-Etched Membrane PCTE filter; GVS Japan, Tokyo, Japan), and 0.7 μm (GF/F glass-fiber filter; Cytiva, Tokyo, Japan).

The on-site filtration is comprised of multiple filter holders connected (PP-47; ADVANTEC, Tokyo, Japan), for which the necessary number of folders were prepared by decontamination through immersion in 10% hypochlorite for at least 30 minutes, followed by adequate washing with water (same for Exp. 2). After filtering the 19:00 samples, 300 mL of ultrapure water was filtered using GF/F filters to serve as filtration-negative controls (Filt-NCs). All filter samples were immediately stored at −20 °C. DNA was extracted from the filter samples in the laboratory, and the mitochondrial cytochrome *b* (cyt*b*) gene and nuclear recombination activating gene 1 (RAG1) of Ayu were quantified using quantitative real-time PCR and species-specific primer-probe sets, respectively (detailed below).

### Experiment 2: Diurnal changes in eDNA concentration during the spawning season

Time-series water sampling was performed between September 20th and 21st, 2021 (the peak of spawning season; https://www.pref.shiga.lg.jp/file/attachment/5387058.pdf; Table S1). 300 mL of surface water was collected in triplicate at 14-time points: 15:00, 17:00, 18:00, 19:00, 20:00, 21:00, 22:00 and 24:00 (day 1), 3:00, 6:00, 9:00, 12:00, 15:00 and 18:00 (day 2). Sampling time points were set to be hourly during the spawning time window from sunset to earlier in the night, and every three hours at other times. The collected water samples were filtered on-site using GF/F filter and filter holder. After filtering all samples, 300 mL of ultrapure water was filtered using GF/F filters as Filt-NCs. All filter samples were immediately stored at −20 °C. DNA was extracted from the filter samples in the laboratory, and the Ayu cyt*b* gene was quantified using a quantitative real-time PCR method with a species-specific primer-probe set (detailed below). However, for the 20:00 and 22:00 samples, Ayu larvae were accidentally captured in 2/3 of the filters, respectively. Thus, these samples were excluded from subsequent analyses. Water and ambient temperatures during the survey were measured at each sampling time point using a bar thermometer and shown in Table S2. From at least five days before the survey date until the end of the survey, the weather was fine, no rain fell and there were no visible changes in water flows. The flow velocity and water depth were measured after water sampling at 18:00, day 2 (Table S3).

### DNA extraction from each filter sample

Firstly, a PCTE filter and a GF/F filter were respectively placed in the lower or upper part of the spin column (EconoSpin, EP-31201; GeneDesign, Inc., Osaka, Japan), with the silica gel membrane eliminated. Prior to DNA extraction, the GF/F filters contained excessive water due to their thickness, which was removed by pre-centrifugation at 5,000 g for 1 minute (Tsuji et al. 2022). Subsequently, 470 μL of a mixture consisting of 200 μL ultrapure water, 250 μL Buffer AL, and 20 μL proteinase K was placed on each filter, and the spin columns were incubated for 45 min at 56 °C. After the incubation, each PCTE filter was re-placed in the upper part of the spin column. The spin columns were then centrifuged at 6,000 g for 1 minute. After adding 500 μL ethanol to the collected liquid and mixing well by pipetting, the DNA solution was transferred to a DNeasy mini spin column (Qiagen, Hilden, Germany). The solution was then purified following the manufacturer’s protocol. Finally, the DNA was eluted in 100 μL of Buffer AE.

### Species-specific primer/probe set development targeting for RAG1 of Ayu

The RAG1 sequences of Ayu and its closely related *Osmeriformes, Hypomesus nipponensis* and *Salangichthys microdon*, were downloaded from the National Center for Biotechnology Information (NCBI) database (https://www.ncbi.nlm.nih.gov/). The species-specific primers and probe were manually designed based on Ayu sequences, where the designed primers have the species-specific nucleotide at the 3′ ends. For the forward primer, the 3′ end is consistent with *S. microdon*; however, there are base mismatches at positions 4, 7 and 19. For the reverse primer has three consecutive mismatches from the 3′ end, so it was judged not to anneal for *S. microdon* DNA (Fig. S3). The TaqMan probe was designed on the amplification range of primers, where it has four and five bases mismatches with *H. nipponensis* and *S. microdon*, respectively. To check the primer parameters and species specificity, *in silico* test was conducted using Primer-BLAST (https://www.ncbi.nlm.nih.gov/tools/primer-blast/) with default settings. Additionally, to check the species specificity of primer/probe set, an *in vitro* test was performed using extracted genomic DNA from Ayu, *H. nipponensis* and *S. microdon* (three individuals for each species). As a DNA template, 100 pg of genomic DNA from each individual was used in each PCR reaction. The real-time PCR conditions were consistent with those used for the analysis of eDNA samples, as described below. Along with assessing the amplification curve by real-time PCR, PCR products were also checked for DNA bands by electrophoresis.

### Quantitative real-time PCR (qPCR) assays

Quantitative real-time PCR was performed in triplicate using two systems (change of equipment due to S.T. transfer): StepOnePlus Real-Time PCR system (Life Technologies, CA, USA; Exp. 1) or Light Cycler 96 system (Roche, Basel, Switzerland; Exp. 2). The species-specific primer-probe set for each target region of Ayu was shown in Table 1 (see Result). All reactions were performed in a 15-μL total volume comprising of 900 nM of each primer (Paa-cyt*b*-F/R or Paa-RAG1-F/R), 125 nM of TaqMan probe (Paa-cyt*b*-Pr or Paa-RAG1-Pr), 0.075 μL AmpErase Uracil N-Glycosylase (Thermo Fisher Scientific, MA, USA), and 2.0 μL DNA template in 1× TaqMan Environmental Master Mix (Thermo Fisher Scientific). In all qPCR runs, a dilution series of linear plasmid DNA at a known pre-set concentration (dilution series of 3 × 10^1^–10^4^ copies per reaction for Exp. 1; 3 × 10^1^–10^5^ copies per reaction for Exp. 2) and PCR negative controls (PCR-NCs, ultrapure water) were also amplified with environmental samples. The linear plasmid DNA was developed by cloning a chimeric sequence of two target sequences of Paa-cytb-F/R and Paa-RAG1-F/R primer into the qTAKN-2 plasmid and digesting it with a restriction enzyme (EcoRI). The qPCR thermal conditions were as follows: 50 °C for 2min, 95 °C for 10 min, followed by 55 cycles of 95°C for 15 s and 60°C for 60 s. For Exp. 1, the R^2^ values for the standard lines of all qPCR for cyt*b* and RAG1 region ranged from 0.996 to 1.0 and 0.996 to 0.999, respectively. For Exp. 2, the R^2^ values for the standard lines of all qPCR for cyt*b* were 1.0. No amplification was observed in any of the Filt-NCs and PCR-NCs.

**Table 1.**
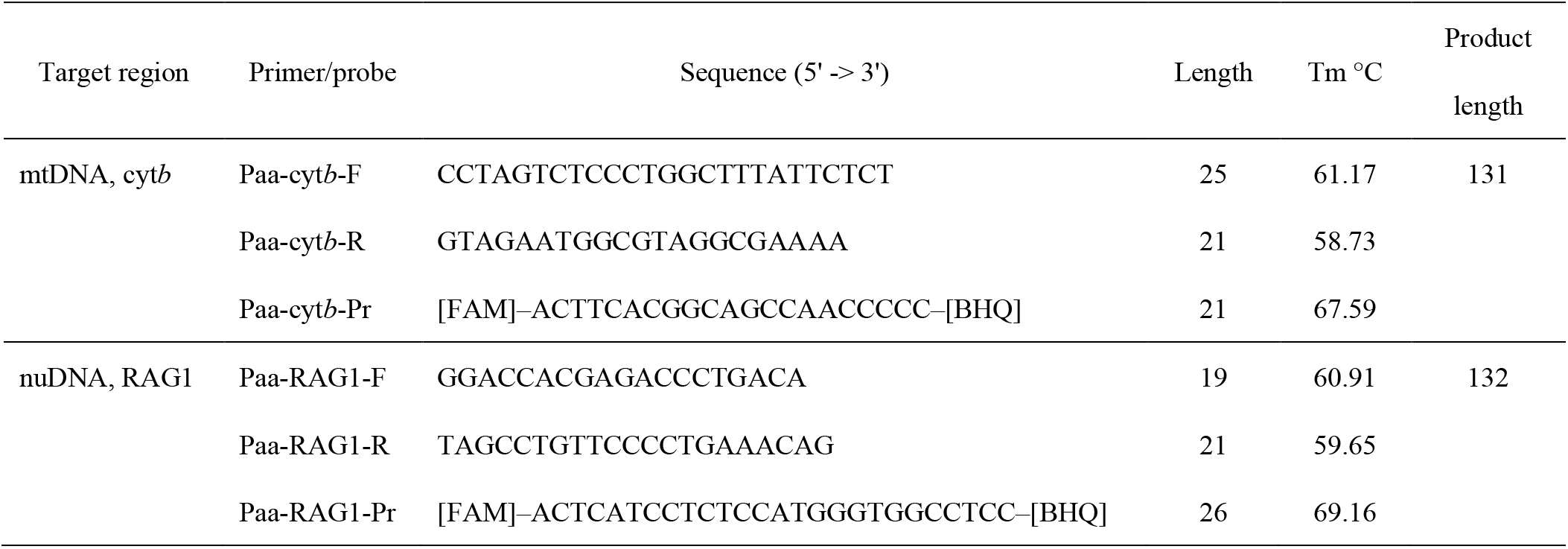
Species-specific primer-probe set for each target region of Ayu, *Plecoglossus altivelis*. Each primer-probe set was developed by Yamanaka & Minamoto (2016) (cyt*b*) and this study (RAG1), respectively.

### Statistical analysis

All statistical analyses for this study were conducted using R version 4.1.1 software (R Core Team. R, 2021). Due to the limited amount of data in Experiment 1 (n = 3), it was difficult to determine the data distribution accurately. Thus, we conducted two statistical analyses, assuming either non-parametric or parametric data distribution in Experiment 1. To examine the overall changes in the cyt*b* or RAG1 concentration and their ratio of eDNA pre- and post-spawning, for each target region, we summed the DNA copies detected on filters of five different pore sizes in each fractional filtration series. We performed the Kruskal–Wallis test followed by the Conover’s test with Holm adjustment (PMCMRplus package; ver. 1.9.3) and one-way analysis of variance (ANOVA) followed by the parametric Tukey honest significant differences (HSD) test. In Figure 2, the p-values obtained from both analyses were shown in lowercase and uppercase, respectively. Additionally, to examine changes in the PSD of eDNA pre- and post-spawning in each sampling day, the concentrations and proportions of eDNA obtained from filters with different pore sizes were compared between daytime and nighttime using the Mann-Whitney U test and t-test for each filter, respectively. To account for the multiplicity of tests, the Bonferroni correction was applied, and we set the minimum level of significance at *p* < 0.01. As the Mann-Whitney U test was considered to have caused Type II errors (see Discussion and Table S5), only the p-values obtained from the t-test are shown in Figure 3.

**Figure 2.**
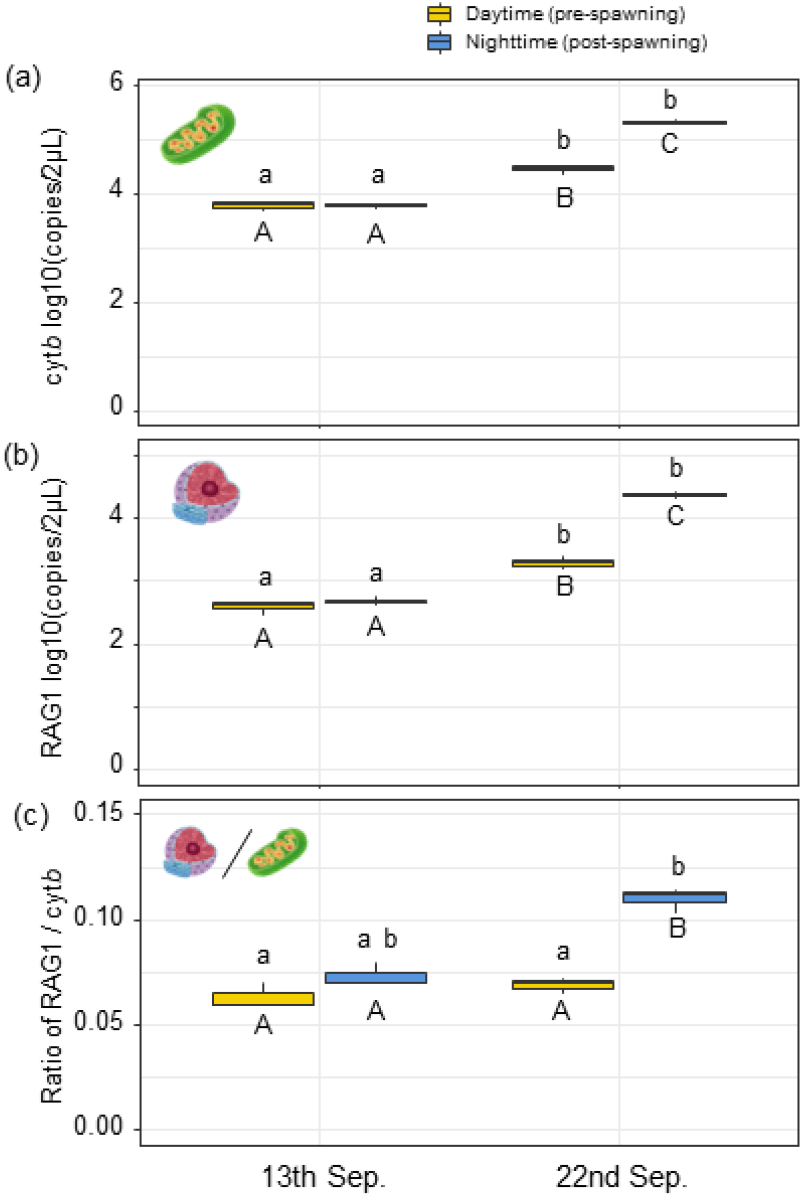
Comparisons of total eDNA concentrations of (a) cyt*b* and (b) RAG1, and (c) nuDNA/mtDNA ratio between daytime (pre-spawning) and nighttime (post-spawning) at two seasons. The orange and blue box plots indicate daytime and nighttime, respectively. The sum of the copy numbers detected on the five filters with different pore sizes was used as the total eDNA concentration. Significant differences are indicated by different letters (lowercase, Conover’s test; uppercase, Tukey HSD test)

**Figure 3.**
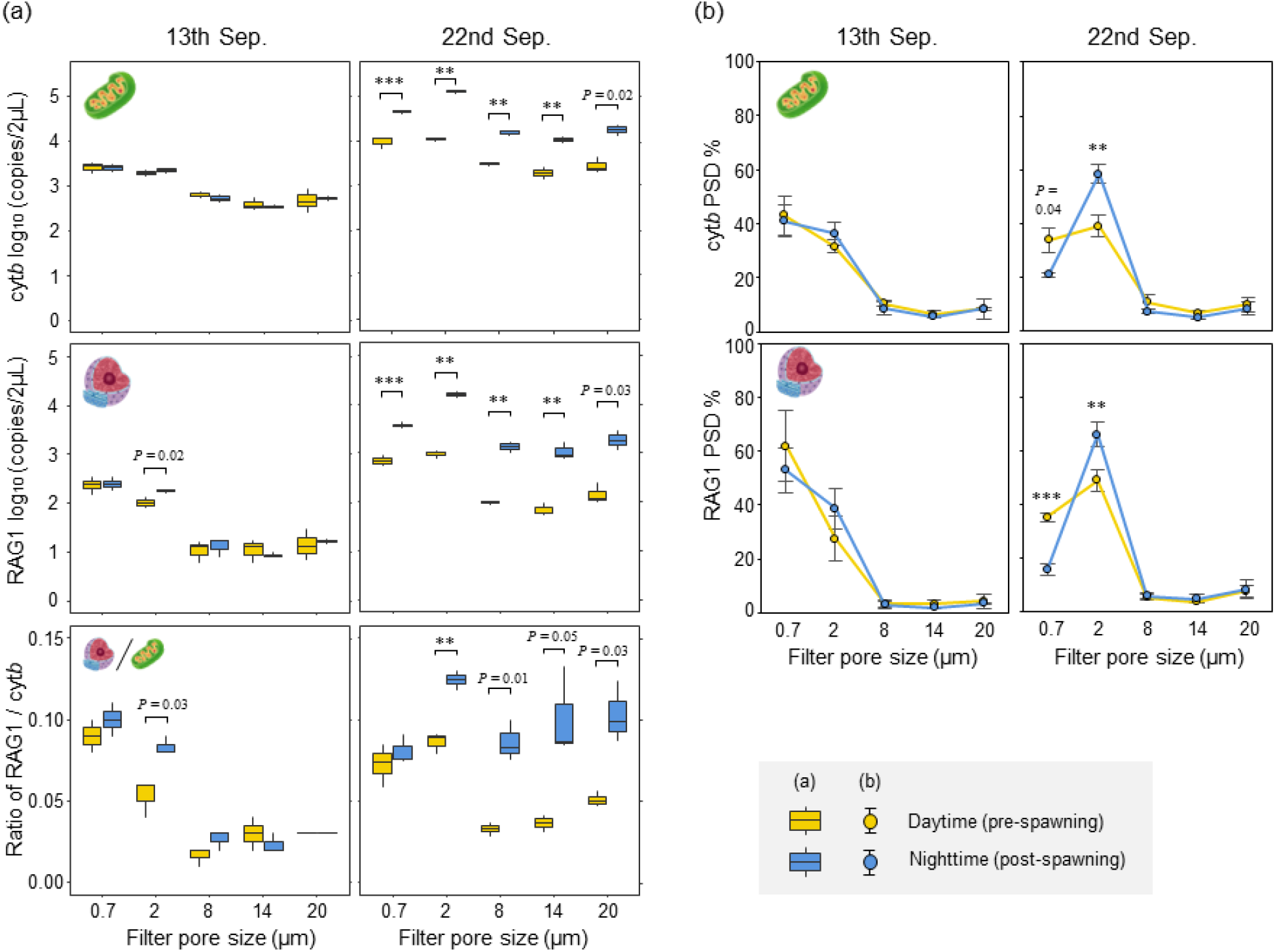
Comparisons of eDNA PSDs and its nuDNA/mtDNA ratio between daytime (pre-spawning) and nighttime (post-spawning) at early (13th Sep.) and peak (22nd Sep.) of spawning seasons. The orange and blue indicate daytime and nighttime. The box plot and the line graph indicate (a) the eDNA concentration and (b) percentage of distribution, respectively. Significant differences are indicated by asterisks (t-test; *p* < 0.01**, *p* < 0.001***; the minimum level of significance was *p* < 0.01)

In Experiment 2, to examine diurnal changes in eDNA concentration, the eDNA concentration observed during the daytime (pre-spawning; 15:00, 17:00, 6:00, 9:00, 12:00, day2-15:00) was compared with those observed at each time point during the nighttime (post-spawning; 18:00, 19:00, 21:00, 24:00, 3:00, day2-18:00) using the Kruskal–Wallis test followed by the Conover’s test with Holm adjustment. The samples, 20:00 and 22:00, were excluded from the analysis because larvae were accidentally trapped on one or two filters out of three replicate filters.

## Results

### Primer probe set development for RAG1 of Ayu

The developed species-specific primers and a probe for the Ayu RAG1 region were shown in Table 1. By *in silico* test using Primer-BLAST, the designed primer-probe set was found to amplify only the target region of Ayu. The estimated primer parameters including primer length, melting temperature (Tm) and product length were almost identical to the previously developed species-specific primer for Ayu cyt*b* region by Yamanaka & Minamoto (2016). Also, in *in vitro* tests using extracted genomic DNA, species-specific amplification was confirmed by real-time PCR and electrophoresis of PCR products.

### Experiment 1: Changes in concentration and PSD of eDNA pre- and post-spawning

The total number of eDNA copies showed a tendency to increase in both DNA regions from the 13th of September towards the peak of spawning on the 22nd (Fig. 2a,b; Table S4). In both DNA regions, a significant increase in total DNA copy number was observed only when comparing pre- to post-sunset on 22 September by Tukey HSD test (*p* < 0.001, both DNA regions; Fig. 2a,b; Table S4). The RAG1 to cyt*b* ratio also showed a significant increase before and after the spawning time window on only 22nd September (Conover’s test, *p* < 0.05; Tukey HSD test, *p* < 0.001; Fig. 2c; Table S4).

On 13th September (early spawning season), even though the obtained P-value did not reach the level of statistical significance, the RAG1 eDNA concentration and nuDNA/mtDNA ratio observed in the 2-8 μm size fraction showed a suggestive increase between pre- and post-spawning (t-test; Fig. 3a; Table S5). On the 22nd of September (the peak of the spawning season), significant increases in eDNA concentrations in both regions were observed at nighttime except for the >20 μm size fraction (t-test; Fig. 3a; Table S5). This increase in eDNA concentration was accompanied by alterations in the PSD, characterised by a decrease in 0.7-2 μm size and an increase in 2-8 μm size post-spawning (t-test; Fig. 3b; Table S5).

### Experiment 2: Diurnal changes in eDNA concentrations during spawning season

During the peak spawning season, a significant change in cyt*b* eDNA concentration was observed, with the highest concentrations occurring within the spawning time window (20:00, 623,250 copies/2μL), approximately 20.5 times greater than the levels observed during the daytime (day1-15:00, 17:00, 6:00, 9:00, 12:00, day2-15:00; average, 30,332 copies/2μL; median, 24,040; SD 19,305 copies/2μL; Fig. 4). The eDNA concentration increased immediately after sunset when Ayu began spawning (Conover’s test, *p* < 0.001), and reached a maximum at 20:00, after which they decreased to the same level as the diurnal concentration seven hours later on 3:00 (Conover’s test, *p* = 1.0). Additionally, on day2-18:00, eDNA concentrations tended to increase again, surpassing the levels observed during the day (Conover’s test, *p* < 0.05).

**Figure 4.**
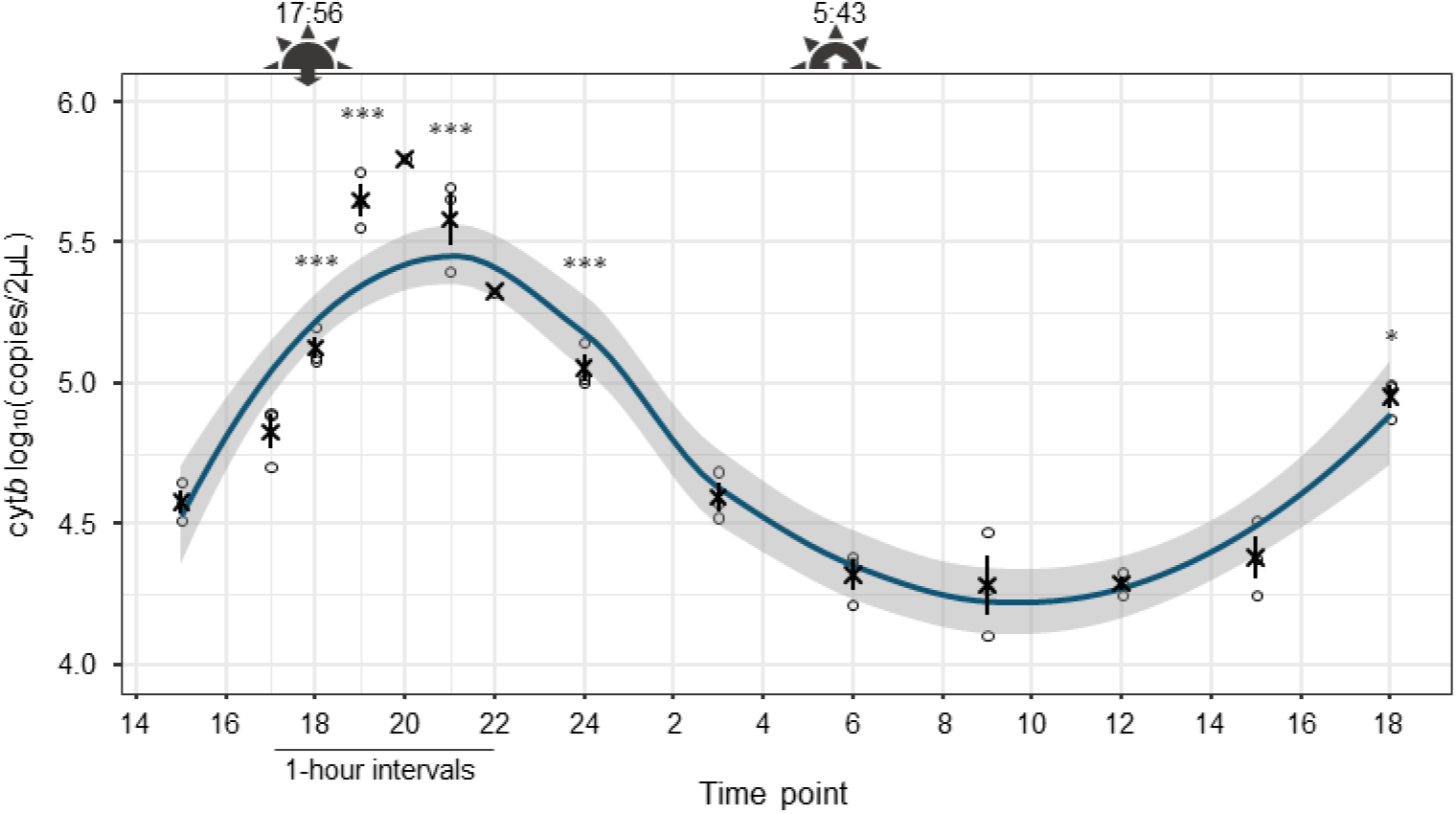
Diurnal changes in eDNA concentration of Ayu cyt*b* during the peak of spawning season. The blue line and grey area indicate a loess regression and 95% CI. The x-mark and bar indicate the mean value and standard error for each time point. Samples at 20:00 and 22:00 were excluded from the analysis due to the accidental capture of Ayu larvae in 2/3 of the filters, respectively. the eDNA concentration observed during the daytime (day 1-15:00, 17:00, 6:00, 9:00, 12:00, day2-15:00) was compared with those observed at each time point during the nighttime by Conover’s test. Significant differences are indicated by asterisks (*p* < 0.05*, *p* < 0.001***). Sunset and sunrise times are as follows: 17:56 and 5:43 Figure SX

## Discussion

In the present study, we have shown that eDNA of the size associated with sperm heads increases post-spawning, and eDNA concentrations had significant diurnal changes with a peak during the spawning time window. Our findings provide essential fundamental information for improving the detection sensitivity and accuracy of eDNA-based spawning surveys. There is no doubt that eDNA analysis can be one of the most efficient and non-invasive strategies for spawning surveys, although attempts to apply it are still in their infancy. To maximise the applicability and usefulness of eDNA analysis for spawning surveys, it is crucial to advance the understanding and accumulation of basic information on the characteristics and dynamics of sperm-derived eDNA in various taxa and habitats in future research.

The elevated eDNA concentrations and nuDNA/mtDNA ratios observed post-spawning at the peak of the spawning season indicate that both can be used as indicators of spawning event occurrence. These results are consistent with previous studies and suggest the high application potential of eDNA analysis for fish spawning surveys (Bylemans et al., 2017; Saito et al., 2022; Tsuji & Shibata, 2021; Wu et al., 2023). Although the non-parametric method did not show statistically significant differences, a trend towards increasing concentrations is evident from Figure 2. This discrepancy should be attributed to the lower statistical power due to the small sample size. On the other hand, our results also suggested that the simple comparison of eDNA concentration and/or the ratio between pre- and post-spawning might not be sensitive enough to identify slight spawning. Although it is estimated that around 0.01% of spawning occurred on 13 September compared to the peak (Table S1), we have missed this slight spawning by both comparisons (Fig. 2). This concern about the sensitivity of eDNA-based monitoring for detecting low-level spawning has also been raised in a previous study on Macquarie perch (*Macquaria australasica*) (Bylemans et al., 2017). As many fish species, including Ayu, aggregate in the spawning site (Azuma, 1973), the concentration of eDNA originating from non-gametes sources (i.e. skin, feces, and mucus etc.) in the water is expected to increase from just before the spawning season to the peak. This baseline increase in eDNA concentration due to fish aggregation might have masked the increase in concentration and ratio due to sperm-derived eDNA released by slight spawning in the early spawning season. Therefore, when aiming to identify a low-level spawning, an analysis strategy that improves detection sensitivity would be necessary.

The characteristic PSD changes observed pre- and post-spawning, i.e. a decrease in 0.7-2 μm and an increase in 2-8 μm (Fig. 3b), would reasonably be explained by sperm size and morphology. The increase in concentration and nuDNA/mtDNA ratio observed pre- and post-spawning is mainly due to the presence of sperm-derived eDNA (Bylemans et al., 2017; Tsuji & Shibata, 2021). The spermatozoa of Ayu is approximately 23 μm in length and consist of a head part containing the nucleus (about 2 μm) and mitochondria (about 0.7 μm), along with a tail part (Fig. S2) (Azuma, 1973; Gwo et al., 1994; Hara, 2009; Hara & Okiyama, 1998). Considering the overall size, sperm-derived eDNA may be trapped in a filter with a pore size of 20 μm. Nevertheless, the increase in eDNA trapped in the 2-8 μm size fraction suggests that the elongated tail of sperm did not prevent passage through the filter and was mainly trapped by the filter depending on the size of their head part. The dependence of PSD of sperm-derived eDNA on head size is consistent with the only previous study using Japanese jack mackerel (*Trachurus japonicus*) (Tsuji et al., 2022). The shapes and sizes of sperm vary widely between species and taxa (Hara & Okiyama, 1998; Kuramoto, 1998). Therefore, comprehending the relationship between more diverse sperm shapes and PSDs in future studies will contribute to a more profound understanding of the eDNA dynamics specific to the spawning period.

Our observations suggest that differences in particle size can be used to semi-selectively recover sperm-derived eDNA, which could be an analytical strategy to improve the sensitivity of eDNA-based spawning detection. Although no significant differences were detected due to the limited power of the statistical analysis due to the lack of samples (n = 3), it is worth noting the clear increasing trend in concentration and ratio post-spawning in the 2-8 μm fraction of the early spawning season that can be read from the graph (*p* = 0.03; Fig. 3a). This result suggests that the increase in eDNA concentration and ratio due to sperm-derived DNA, which were masked by tissue-derived DNA when all sizes of eDNA were considered, would be observed more clearly by the fractional recovery. Fractional filtration is undeniably slightly more labour-intensive than normal filtration, but in practical terms, this is unlikely to have a serious impact on feasibility because we are only required to use two filters with a pore size that takes into account the sperm head size. Accurate and non-invasive detection of entry into the spawning season using eDNA analysis is expected to contribute to the understanding of spawning ecology and resource management.

The significant diurnal changes in eDNA concentrations observed during the spawning season would reflect the spawning time window of Ayu. Ayu is known to spawn in the gravel beds in riffles every day during the spawning season from around sunset to a few hours (Miyadi, 1960; Takahashi & Azuma, 2016), and it was consistent with our results. Diurnal changes in fish eDNA concentrations could be caused by simple changes in biomass or activity levels as well as spawning, but previous studies and our observations would rule out these possibilities. In the only previous study about diurnal changes in eDNA concentration, a slight increase in eDNA levels associated with nocturnal activity in nocturnal fish, Japanese eels (*Anguilla japonica*), was reported by aquarium experiments (Takahashi et al., 2021). In contrast, previous studies on nocturnal amphibians found no differences in eDNA concentrations between daytime and nighttime (*Cryptobranchus a. alleganiensis*, Takahashi et al., 2018; *Ascaphus montanus* and *Dicamptodon aterrimus*, Pilliod et al., 2013). Ayu, the target species in this study, is a diurnal fish, and their activity level at nighttime is significantly lower than during the daytime in the non-spawning season (Minh-Nyo et al., 1991; Miyadi, 1960). Additionally, Ayu aggregates in the lower reaches of rivers in the spawning season, but they do not actively migrate during the day and night and stay in slowly flowing areas near the spawning sites (Takahashi & Azuma, 2016). Additionally, the fact that no differences in eDNA concentrations were observed before and after sunset during the early spawning season (13th September) makes it reasonable to assume that Ayu eDNA does not usually show significant diurnal changes with peaks at night. Therefore, the substantial diurnal changes in eDNA concentrations observed in this study were caused by spawning behaviour and are expected to be specific to the spawning season. This is the first study to monitor and report diurnal changes in eDNA concentrations during the spawning season.

The presence of diurnal changes in eDNA concentration has implications for the planning of sampling design in future eDNA surveys. Firstly, in spawning surveys based on eDNA, the timing of sampling should be determined by whether the researcher wants to estimate the occurrence and amounts of spawning or fish aggregation amounts. In previous spawning surveys based on eDNA analysis, the spawning time has not been considered in determining the timing of eDNA sampling. Almost all previous results from eDNA-based spawning surveys conducted in natural habitats showed a reasonable relationship with the abundance of aggregated fish observed using conventional methods, but either contradicted or showed very weak relationships with the abundance of eggs and larval recovered (Bylemans et al., 2017; Erickson et al., 2016; Hayer et al., 2020; Inui et al., 2021; Thalinger et al., 2019; Tillotson et al., 2018; Yatsuyanagi et al., 2020). This may be because sperm-derived eDNA is released and subsequently diffuses downstream, even if the fish remain in the vicinity of the spawning area, with concentrations changing significantly during several hours. So, in estimating the occurrence and/or amounts of spawning, it would be detected more sensitively and accurately by sampling before and after the spawning time window and comparing concentrations and ratios. On the other hand, when estimating fish aggregation amounts, sampling as long as possible after the spawning time window would reduce the effect of increased concentration due to sperm-derived eDNA. However, it is assumed that diurnal changes in eDNA concentrations will not be observed or will be weakened in species that spawn regardless of time and in lentic environments. Thus, it would be desirable to investigate the relationship between spawning and dynamics of eDNA concentration in species and habitats with diverse spawning ecology in the future and to continue to seek further sampling strategies to appropriate research objectives.

Sampling focused on the spawning time window during the spawning season is likely to benefit not only the spawning survey but also the presence/absence assessments of species with extremely small biomass, such as invasive species in the early established stage and endangered species. The low eDNA concentration due to the small biomass is one of the main causes of false-negative results in eDNA studies (Carim et al., 2019; Jerde et al., 2013). Thus, some previous studies have suggested that eDNA sampling during the spawning season, when eDNA concentrations are likely to increase for species with small biomass, may be effective in avoiding false negative results (Bracken et al., 2019; Crane et al., 2021; Tsuji & Shibata, 2021). Our results also support this view and it seems reasonable to focus sampling on the spawning season, when the baseline of eDNA concentrations increases. Furthermore, we suggest sampling during the spawning time window when the eDNA concentrations are at their peak because sperm-derived eDNA is subject to diffusion and degradation. This sampling strategy should further improve detection sensitivity. Increased detection sensitivity will contribute to expanding the applicability of eDNA analysis and leading to reliable results.

## Acknowledgements

We thank Dr. Hideyuki Doi (Kyoto university) for his accurate and valuable advice on the manuscript. All experiments in the present study were compliant with the current laws of the country, Japan, in which they were performed.

## Statements and Declarations

We declare no conflicts of interest.

## Availability of data and material

Full details of the qPCR results for each experiment of the present study are available in the supporting information (Table S6 and S7).

## Authors’ contributions

S.T.; Conceptualization, Methodology, Fieldwork (Exp. 2), Molecular analysis, Visualization, Writing − original draft, Funding acquisition: N.S.; Fieldwork (Exp. 1 and 2), Molecular analysis, Writing − review & editing.

## Figure SX

**Figure S1.**
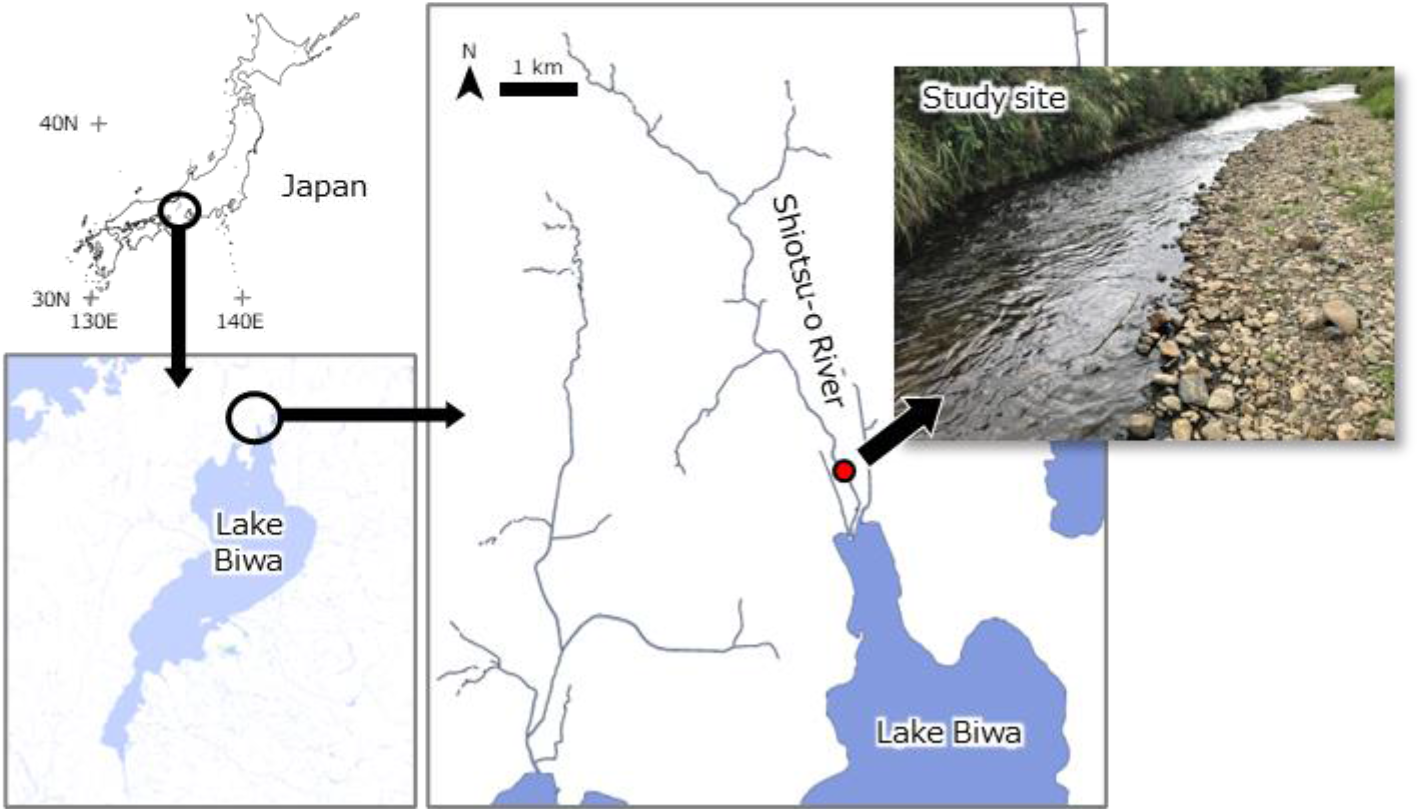
Study site.

**Figure S2.**
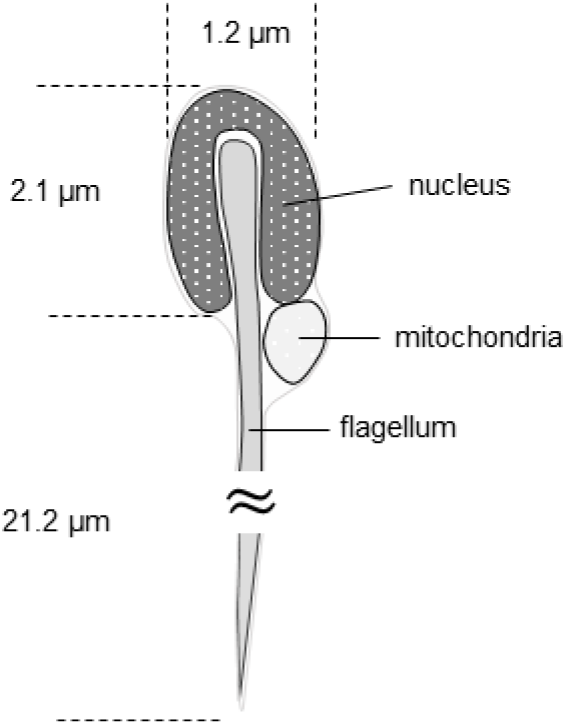
Simple schematic representation of the ultrastructure of Ayu, P. altivelis, sperm.

**Figure S3.**
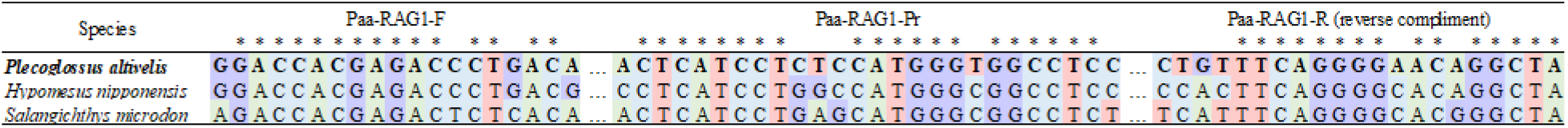
Sequences of the target region of the designed primer-probe set for nuclear recombination activating gene 1 (RAG1) of Ayu, P. altivelis, and its closely related species in Japan. The amplicon length was 132 bp.

